# Ecological conditions experienced by bat reservoir hosts predict the intensity of Hendra virus excretion over space and time

**DOI:** 10.1101/2021.08.19.457011

**Authors:** Daniel J. Becker, Peggy Eby, Wyatt Madden, Alison J. Peel, Raina K. Plowright

## Abstract

The ecological conditions experienced by wildlife reservoir hosts affect the amount of pathogen they excrete into the environment. This then shapes pathogen pressure, the amount of pathogen available to recipient hosts over space and time, which affects spillover risk. Few systems have data on both long-term ecological conditions and pathogen pressure, yet such data are critical for advancing our mechanistic understanding of ecological drivers of spillover risk. To identify these ecological drivers, we here reanalyze shedding data from a spatially replicated, multi-year study of Hendra virus excretion from Australian flying foxes in light of 25 years of long-term data on changing ecology of the bat reservoir hosts. Using generalized additive mixed models, we show that winter virus shedding pulses, previously considered idiosyncratic, are most pronounced after recent food shortages and in bat populations that have been displaced to novel habitats. We next derive the area under each annual shedding curve (representing cumulative virus excretion) and show that pathogen pressure is also affected by the ecological conditions experienced by bat populations. Finally, we illustrate that pathogen pressure positively predicts observed spillover frequency. Our study suggests that recent ecological conditions of flying fox hosts are shifting the timing, magnitude, and cumulative intensity of Hendra virus shedding in ways that shape the landscape of spillover risk. This work provides a mechanistic approach to understanding and estimating risk of spillover from reservoir hosts in complex ecological systems and emphasizes the importance of host ecological context in identifying the determinants of pathogen shedding.

## Introduction

Cross-species transmission can be conceptualized as a series of hierarchical barriers that must be overcome for a pathogen to move from a reservoir host to recipient hosts such as humans [1]. At the beginning of this cascade, the ecological conditions experienced by reservoir hosts shape susceptibility to infection and the likelihood of pathogen shedding (i.e., excretion) [2,3]. For example, droughts have been associated with increased nematode shedding in ungulates [4], and *Salmonella* shedding by ibis is more likely in urban than rural habitats [5]. This in turn determines pathogen pressure, or the amount of pathogen available to recipient hosts over space and time, which affects the force of infection and the ultimate probability of spillover [6]. Identifying the ecological conditions that drive pathogen pressure could accordingly improve reservoir host surveillance efforts and help predict where and when spillover is most likely [7,8].

Multiple challenges restrict our ability to establish the ecological drivers of pathogen pressure. On the one hand, many host populations must be repeatedly sampled to capture spatial and temporal variation in shedding [9]. This presents logistical difficulties and often necessitates optimized allocation of sampling effort [10,11]. On the other hand, the complex ecological dynamics used as predictors are best characterized with long-term research [12,13]. For example, more than two decades of annual monitoring were necessary to identify the ecological conditions that predict the density of infected nymphal ticks and human Lyme disease risk [14]. As such, the general sparsity of long-term, ecological data in many wildlife pathogen systems limits inference about the possible factors that shape pathogen pressure from reservoir hosts. Systems with data on pathogen pressure and long-term ecological conditions are critical for advancing our mechanistic understanding of spillover and to identify likely drivers in less studied systems.

Hendra virus (HeV) shedding from Australian flying foxes (*Pteropus* spp.) provides an ideal system to identify the ecological conditions that predict pathogen pressure [15]. HeV emerged in 1994, causing an outbreak of a lethal disease in horses and then humans that has been followed by over 60 spillovers through 2019 across eastern Australia [16,17]. Flying foxes are the natural reservoirs of HeV [18], and transmission to horses likely occurs via contact with pasture, feed, or water contaminated with urine, saliva, or feces [19]. Spatiotemporal sampling of bat roosts has revealed shedding pulses during the Austral winter in subtropical regions, yet their occurrence varies both spatially and interannually [20]. This suggests that seasonal processes such as births and climate shifts cannot solely drive pathogen pressure [21]. The mechanisms underlying shedding pulses remain poorly understood but have been hypothesized to stem from synchronous ecological conditions that affect bat behavior and within-host processes [22–25].

Traditionally, flying foxes move nomadically to find ephemeral sources of nectar in native forests [26,27]. However, few dietary plants reliably flower in the Austral winter [27,28]. These plants occur in forests that have been mostly cleared for agriculture and urbanization, making native winter feeding habitat scarcer and more dispersed [29]. Eucalypt flowering is especially vulnerable to disruption from temperature and rainfall variability [28,30], and occasionally winter or spring nectar fails, leading to acute food shortages and periods of nutritional stress [27,31,32]. Bats have responded to these changes by splitting into smaller roosts that are closer to reliable, often introduced, food in agricultural areas and urban gardens [27,33]. Bats in these newly established and continually occupied roosts supplement their diet with non-native plants [34,35]. This change in behavior has expanded the winter distribution of flying foxes in subtropical Australia into areas that do not provide native winter food [36,37].

Reliance on agricultural and urban dietary resources when native plants are unavailable could combine with other energetic stressors, such as seasonal reproduction and the more extreme temperatures that bats can experience in these newly established roosts outside their historic wintering range [38–40]. These stressors could alter bat within-host dynamics (e.g., immunosuppression) in ways that increase susceptibility to HeV or facilitate reactivation from chronic infections [22,24,25,41]. Periods of nutritional stress have been associated with poor body condition and elevated HeV seroprevalence in little red flying foxes (*P. scapulatus*) [31]; moreover, poor condition was a correlate of HeV infection in black flying foxes (*P. alecto*) but was negatively correlated with HeV seroprevalence in *P. alecto* and grey-headed flying foxes (*P. poliocephalus*) [42,43]. The relationships between food shortages and HeV shedding therefore remain poorly understood, especially in the context of the changing ecological conditions experienced by these bat hosts. Recent work suggests such ecological changes have increased the frequency of HeV spillovers (Eby et al., in review) [17,44], but the mechanisms linking food shortages and dramatic shifts in bat behavior with virus shedding dynamics remain unresolved.

Here, we reanalyze shedding data from a spatially replicated, multi-year study of HeV in flying foxes in light of long-term data on bat ecology to test whether food shortages and recent shifts in flying fox behavior amplify pathogen pressure. We first ask whether these ecological conditions can explain seemingly idiosyncratic seasonal shedding pulses while also accounting for underlying spatiotemporal variation in infection dynamics and roost composition, including species with high competence (e.g., *P. alecto*) [45–47]. We next quantify pathogen pressure per year and per roost and test whether bat ecology also predicts cumulative seasonal shedding intensity across the landscape. Finally, we assess the relationship between pathogen pressure and the frequency of realized spillover events. Our work thus provides a mechanistic approach to understanding bat-borne virus spillover risk as well as a new method for estimating these risks from multiple ecological predictors. Bats throughout the eastern hemisphere are experiencing similar stressors owing to land conversion [23,48], underscoring the general need to better understand the likely consequences for pathogen shedding and subsequent spillover risk.

## Materials and methods

### Spatiotemporal data on HeV shedding

We reanalyzed a previously published longitudinal dataset of HeV shedding from flying foxes spanning July 2011 to November 2014 across New South Wales and Queensland [20]. Sampling consisted of urine collection from quadrants of 3.6 m x 2.6 m plastic sheets placed under roosts at primarily monthly intervals [49]. Urine was pooled per quadrant and screened by quantitative RT-PCR targeting the HeV M gene to determine presence or absence of viral shedding in a roost [50]. We aggregated data to the scale of week and determined HeV prevalence as the proportion of positive urine pools. Similarly, we calculated the median number of all flying foxes and for each Australian *Pteropus* species per roost per time interval. We restricted HeV data to an area of subtropical eastern Australia (i.e., mid-to-northeast New South Wales, southeast Queensland) that represents our study region for long-term data on bat ecology (see below) and includes the locations of almost all subtropical HeV spillovers [15]. We further restricted analyses to 2012 through 2014, as 2011 sampling mostly occurred at a single roost [20]. We also limited analyses to roosts sampled in at least two of the three years and in at least three timepoints per year. We thus analyzed prevalence data from nine roosts sampled in 2012–2014 (*n*=196 weeks; Fig. 1).

**Figure 1.**
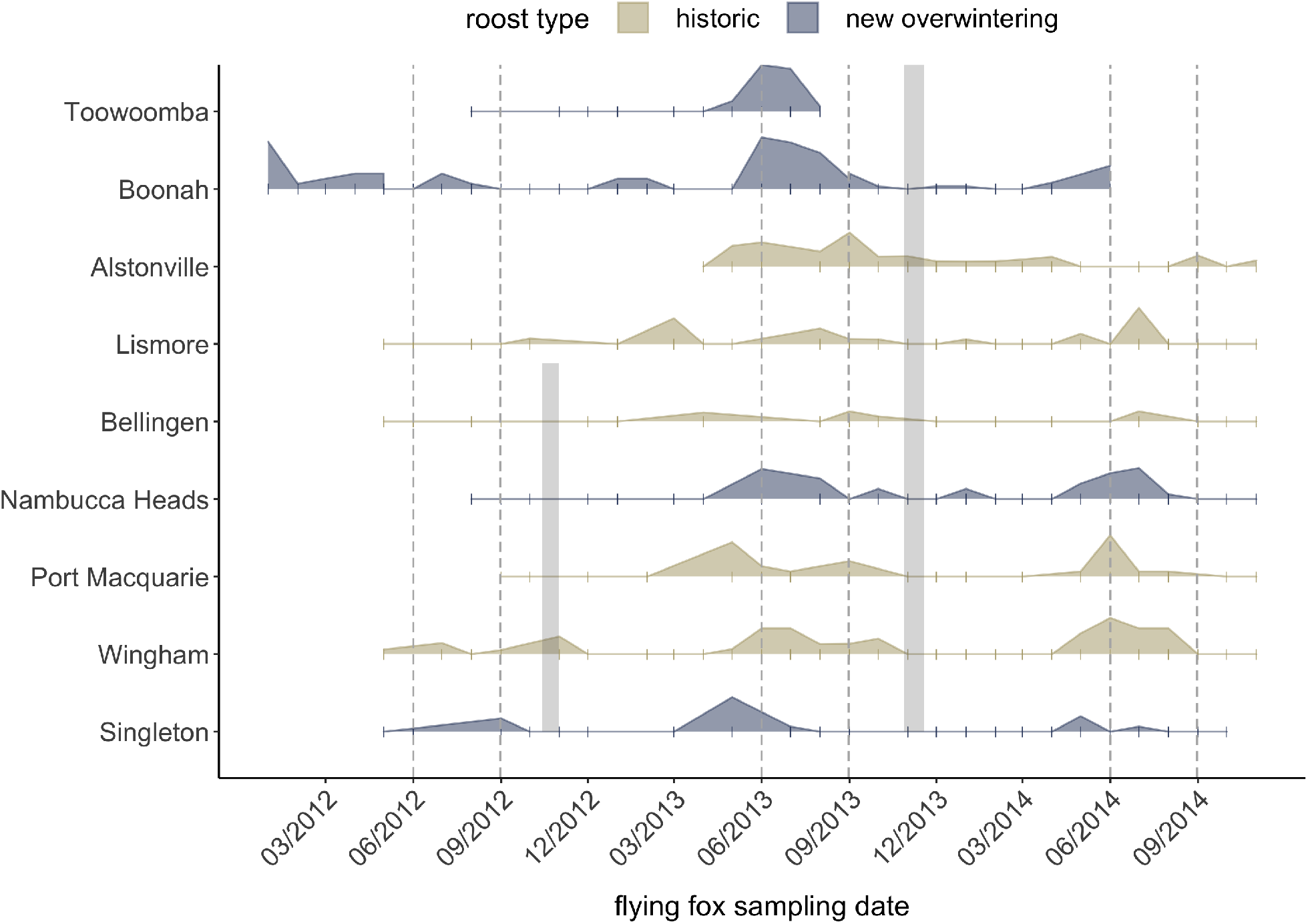
Spatiotemporal variation in HeV shedding for the nine flying fox roosts. Curves display the weekly proportion of HeV-positive urine pools, with roosts shown in order of latitude and colored by roost type. Ticks show sampling timepoints. Grey shading indicates regional acute food shortage events, and dashed lines indicate the Austral winter (i.e., June through August).

### Roost-level ecological conditions

We paired HeV prevalence data within our study region with long-term data on flying fox behavior. As described in full in Eby et al. (in review), we used data held by Australian state governments, flying fox surveys conducted between 1998–2005 and 2012–2019, records held by local land managers and experienced observers, and direct observation by the authors to characterize the ecological conditions experienced by bat populations over time [27,28,37]. Roosts were classified as belonging to their historic winter range (and thus where bats rely on native winter dietary plants) or to newly established overwintering regions (and thus where native food is unavailable). We also used these data to identify regional food shortages, acute periods associated with reduced flowering of native dietary plants or no flowering (Table 1).

**Table 1.** Spatiotemporal ecological conditions of flying fox roosts included in the analysis.

| Roost | Longitude | Latitude | Year | Food shortage in prior year | Roost type |
| --- | --- | --- | --- | --- | --- |
| Toowoomba | 151.954 | -27.564 | 2012 | 0 | New overwintering |
|  |  |  | 2013 | 0 |  |
| Boonah | 152.681 | -27.992 | 2012 | 0 | New overwintering |
|  |  |  | 2013 | 0 |  |
|  |  |  | 2014 | 1 |  |
| Alstonville | 153.439 | -28.481 | 2013 | 0 | Historic |
|  |  |  | 2014 | 1 |  |
| Lismore | 153.277 | -28.807 | 2012 | 0 | Historic |
|  |  |  | 2013 | 0 |  |
|  |  |  | 2014 | 1 |  |
| Bellingen | 152.897 | -30.452 | 2012 | 0 | Historic |
|  |  |  | 2013 | 1 |  |
|  |  |  | 2014 | 1 |  |
| Nambucca Heads | 153.003 | -30.642 | 2012 | 0 | New overwintering |
|  |  |  | 2013 | 1 |  |
|  |  |  | 2014 | 1 |  |
| Port Macquarie | 152.908 | -31.431 | 2012 | 0 | Historic |
|  |  |  | 2013 | 1 |  |
|  |  |  | 2014 | 1 |  |
| Wingham | 152.376 | -31.871 | 2012 | 0 | Historic |
|  |  |  | 2013 | 1 |  |
|  |  |  | 2014 | 1 |  |
| Singleton | 151.175 | -32.562 | 2012 | 0 | New overwintering |
|  |  |  | 2013 | 1 |  |
|  |  |  | 2014 | 1 |  |

### Analyses of HeV shedding

To test if roost type (i.e., historic or new overwintering regions) and regional food shortages affect seasonal HeV shedding, we fit generalized additive mixed models (GAMMs) with a binomial response to urine pool prevalence with the *mgcv* package in R [51,52]. GAMMs can flexibly approximate the temporal dynamics of infectious diseases, even when underlying transmission mechanisms are unknown [53,54]. Our fixed effects included a cyclic cubic spline for week, ordered factors of roost type and regional food shortage, and all possible interactions. We used regional food shortages from the prior year (i.e., October 2012 and October–November 2013; Fig. 1) owing to previously observed associations between spring and summer food shortages and subsequent winter spillovers of HeV in 2010–2011 and in 2016–2017 [44].

We also adjusted for occupancy of *P. alecto* using its relative abundance per roost from weekly median flying fox counts. We did not account for co-roosting *P. poliocephalus* and *P. scapulatus*, as *P. alecto* is the only species that has been associated with HeV shedding in our study region [45,46]. As flying foxes roost in aggregations of up to hundreds of thousands of individuals, counts and resulting species proportions are estimates. We thus derived a binary variable denoting if the weekly proportion of *P. alecto* per roost was below or above 25%, the lowest maximum proportion of this species across all roosts in this dataset (Fig. S1) [20]. This cutoff accordingly indicates the upper occupancy bound of this species when it is rare. Our GAMMs also controlled for spatial dependence through a bivariate smooth of longitude and latitude [55]. To account for residual variation in spatiotemporal shedding, we included a random factor smooth of week per roost per year [56]. We set the random effect such that all groups can differ in weekly shedding but are penalized if they deviate too strongly from the global trend.

Given the complexity of our GAMM relative to our sample size, we used an information theoretic approach to select among competing nested fixed effects. We considered (*i*) our full model alongside two simplified GAMMs: (*ii*) without the three-way interaction between week, roost type, and prior year food shortage and (*iii*) without the two-way interaction between the latter categorical predictors. We fit GAMMs with maximum likelihood (ML) and compared models using corrected Akaike information criterion (AICc) and Akaike weights (*w_i_*). We considered models within two ΔAICc to be competitive [57]. We then refit our most supported models with restricted ML (REML) for parameter estimation and visualizing fitted values.

To quantify cumulative annual shedding intensity, we fit univariate GAMs with binomial response and cyclic cubic splines for week to our HeV prevalence data per year for each of our nine roosts (*n*=25; Fig. S2). In rare cases with few (e.g., *n*=3) sampling events and where prevalence was uniformly zero (i.e., as occurred in two annual time series; Fig S2), we used thin plate splines to improve convergence. We then calculated the area under each annual shedding curve (i.e., pathogen pressure; AUC) by integrating the fitted values and confidence intervals using the *Bolstad2* package [53,58]. This measure summarizes the magnitude and duration of shedding in a single metric, and thus greater pathogen pressure suggests more cumulative HeV-positive pools per roost per year. We used the *metafor* package to assess heterogeneity in AUC through a random-effects model weighted by sampling variance, which we derived from the number of sampling events per time series [59]. We then used REML to quantify *I^2^*, which measures the contribution of true heterogeneity rather than noise to the variance in AUC [60].

We then assessed the ecological drivers of pathogen pressure using another set of GAMs. We modeled annual AUC as a Tweedie-distributed response and again weighted these estimates by the number of sampling events per annual time series. We included roost type and prior food shortage as categorical predictors alongside a bivariate smooth of longitude and latitude. We also controlled for the mean annual proportion of *P. alecto* per roost with a thin plate spline and assessed sensitivity of our results to two alternative measures of annual relative abundance (e.g., median, winter maximum). Lastly, because a small number of our annual time series had short sampling durations that did not include the Austral winter (Fig. S2), we reran our GAMs after excluding four annual AUC estimates derived from 20 or fewer weeks of flying fox sampling.

### HeV spillover analyses

To assess relationships between pathogen pressure and spillover, we collated data on the location and date of all known HeV cases in horses in our study region between 2012–2014 [15]. As described in full in Eby et al. (in review), these data were obtained from government notices (i.e., New South Wales Department of Primary Industries, Queensland Department of Agriculture and Fisheries; Business Queensland), ProMED, local media reports, and personal communications. We aggregated all HeV case locations to the nearest town or regional center for confidentiality.

We derived spatial buffers using the *rgeos* package to collate the total number of HeV spillover events near each of our nine roosts per sampled year. Australian *Pteropus* are highly mobile [32,61], and prior analyses have shown weak spatial synchrony in HeV shedding among flying fox roosts up to approximately 500 km [21]. We therefore used spatial buffers of 50, 100, 200, 300, 400, and 500 km. We modeled counts of spillovers within each buffer through a set of GAMs that used Poisson distributions and penalized splines for pathogen pressure [62]. As in our GAM analyses of AUC, we also included a bivariate smooth of longitude and latitude.

## Results

### Ecological predictors of seasonal HeV shedding pulses

Our GAMMs revealed that previously observed seasonality in HeV shedding [e.g., 26] is markedly explained by recent changes in bat behavior and by food shortages. The most parsimonious model explained 71% of the deviance and only included interactions between week and both roost type and recent food shortages (*w_i_*=0.59; Table S1). Ordered factor smooths indicated that roosts in newly established overwintering regions had distinct seasonal shedding patterns compared to roosts in the historic wintering range (*χ^2^*=23.61, *p*<0.001); roosts that experienced a recent food shortage also displayed distinct shedding seasonality compared to roosts without recent food shortages (*χ^2^*=14.33, *p*<0.001; Table 2). In the absence of recent food scarcity, historic roosts had negligible seasonality in shedding, while new overwintering roosts exhibited shedding pulses in winter (Fig. 2A). Yet when food shortages occurred in the spring prior to sampling (Fig. 2B), both historic and new overwintering roosts displayed stronger shedding seasonality and amplitude the following winter. Further, the winter shedding amplitude for new roosts was greater than historic roosts. Such patterns held after accounting for relative *Pteropus alecto* occupancy (Fig. 2C–D), which positively predicted HeV prevalence (Table 2).

**Table 2.**
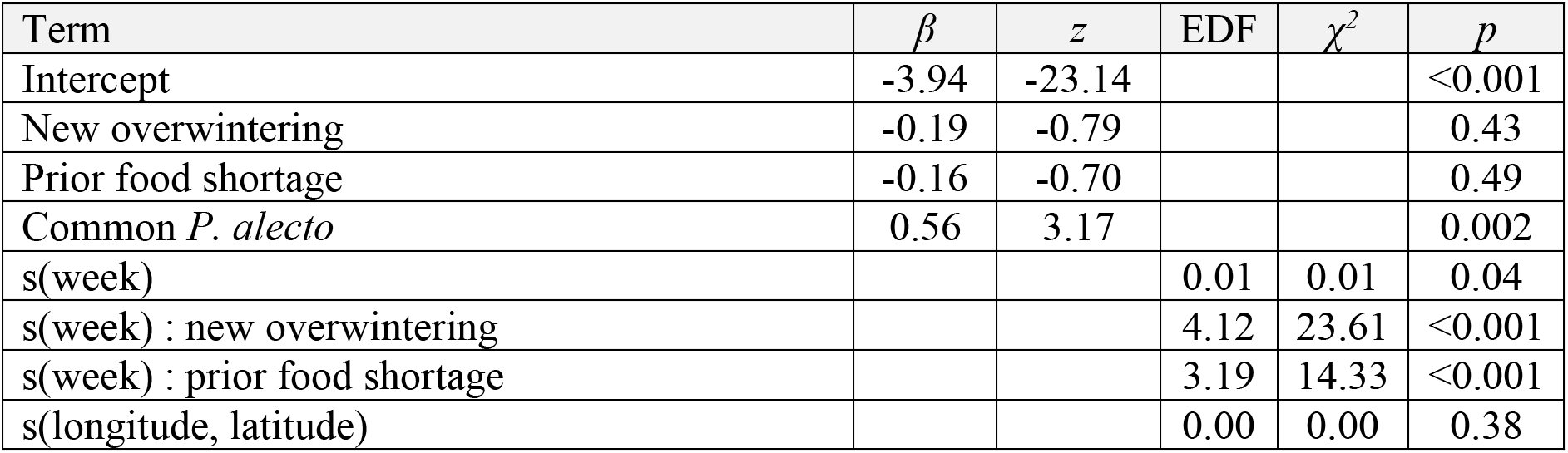
Results of the most parsimonious GAMM for predicting seasonal HeV shedding from Australian flying foxes (*n*=196), fit using REML. Fixed effects are presented as ordered factors with coefficients (categorical) or the estimated degrees of freedom (EDF) and test statistics.

| Term | $\beta$ | $z$ | EDF | $\chi^2$ | $p$ |
| --- | --- | --- | --- | --- | --- |
| Intercept | -3.94 | -23.14 |  |  | <0.001 |
| New overwintering | -0.19 | -0.79 |  |  | 0.43 |
| Prior food shortage | -0.16 | -0.70 |  |  | 0.49 |
| Common *P. alecto* | 0.56 | 3.17 |  |  | 0.002 |
| s(week) |  |  | 0.01 | 0.01 | 0.04 |
| s(week) : new overwintering |  |  | 4.12 | 23.61 | <0.001 |
| s(week) : prior food shortage |  |  | 3.19 | 14.33 | <0.001 |
| s(longitude, latitude) |  |  | 0.00 | 0.00 | 0.38 |

**Figure 2.**
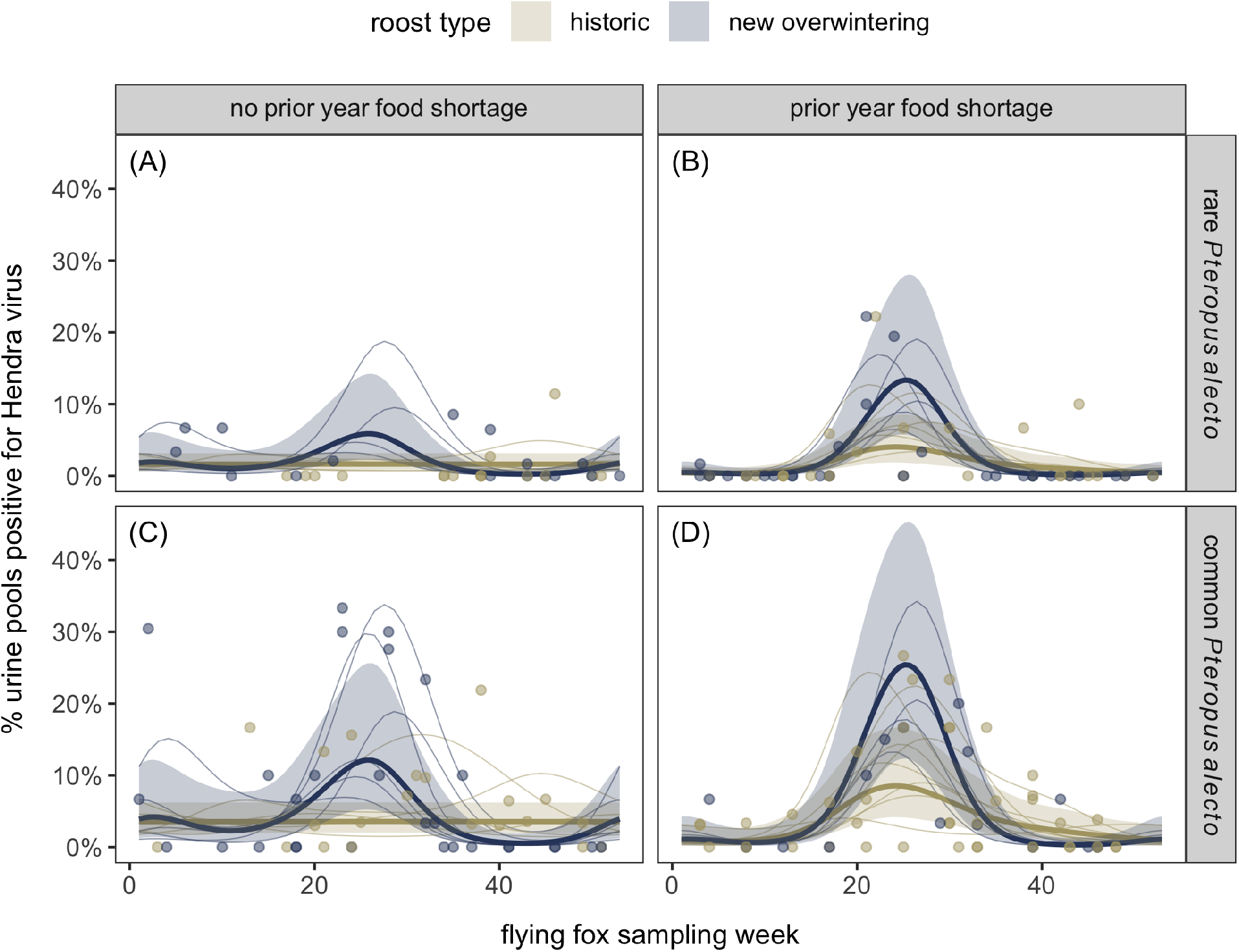
Fitted HeV urine pool prevalence and 95% confidence intervals from the most parsimonious GAMM with week, seasonal interactions with roost type and previous food shortages, the interaction between both factor variables, and an adjustment for categorical weekly *Pteropus alecto* occupancy. Weekly data are overlaid and colored by roost type. Thin lines show the fitted curves from the random factor smooth including each roost per year.

### Heterogeneity in pathogen pressure

Cumulative HeV pathogen pressure (i.e., area under each annual shedding curve) varied substantially across roosts and years (Fig. 3A). Our meta-analysis identified significant heterogeneity among AUC estimates (*I^2^*=0.93, *Q_24_*=339.59, *p*<0.001). When testing the ecological drivers of this variation, our GAM explained 40% of the deviance in pathogen pressure (Table 3). Whereas recent food shortages had little effect on AUC (*β*=0.15, *t*=1.38, *p*=0.19), newly established overwintering roosts had significantly greater pathogen pressure than historic roosts (*β*=0.76, *t*=4.33, *p*<0.001; Fig 3B). Similar results were obtained when adjusting for median annual and maximum winter fractions of *P. alecto* per roost (Table S2 & S3). In our primary model, the mean annual proportions of *P. alecto* also significantly and largely positively predicted AUC (*F*=7.88, *p*<0.001); similar results were obtained with median annual proportions of *P. alecto* but not winter maximum occupancy (Fig. S3; Table S2 & S3). Lastly, excluding the four AUC estimates from truncated time series mostly strengthened the associations between predictors (roost type, food shortage, *P. alecto*) and pathogen pressure (Fig. S4; Table S4–6).

**Table 3.**
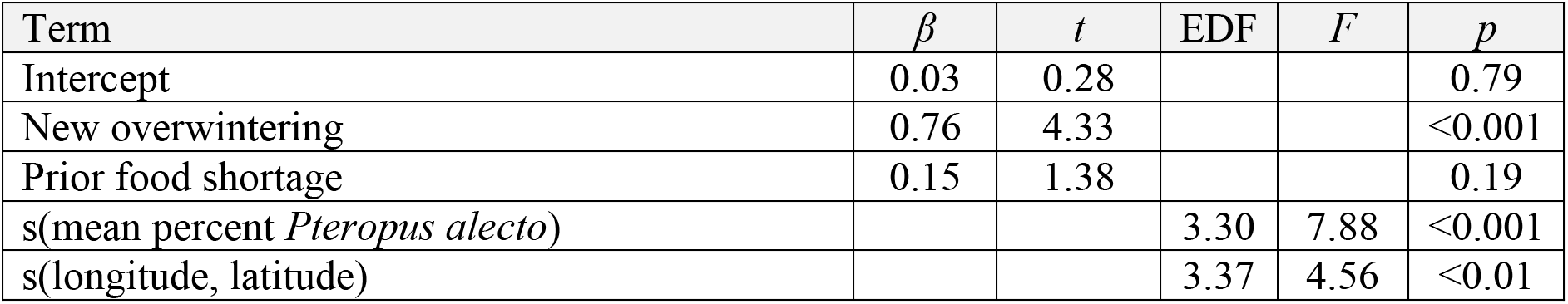
Results from the GAM of annual AUC estimates from flying fox roosts (*n*=25), fit using REML. Fixed effects are presented as ordered factors with model coefficients (categorical) or the estimated degrees of freedom (EDF) and test statistics.

**Figure 3.**
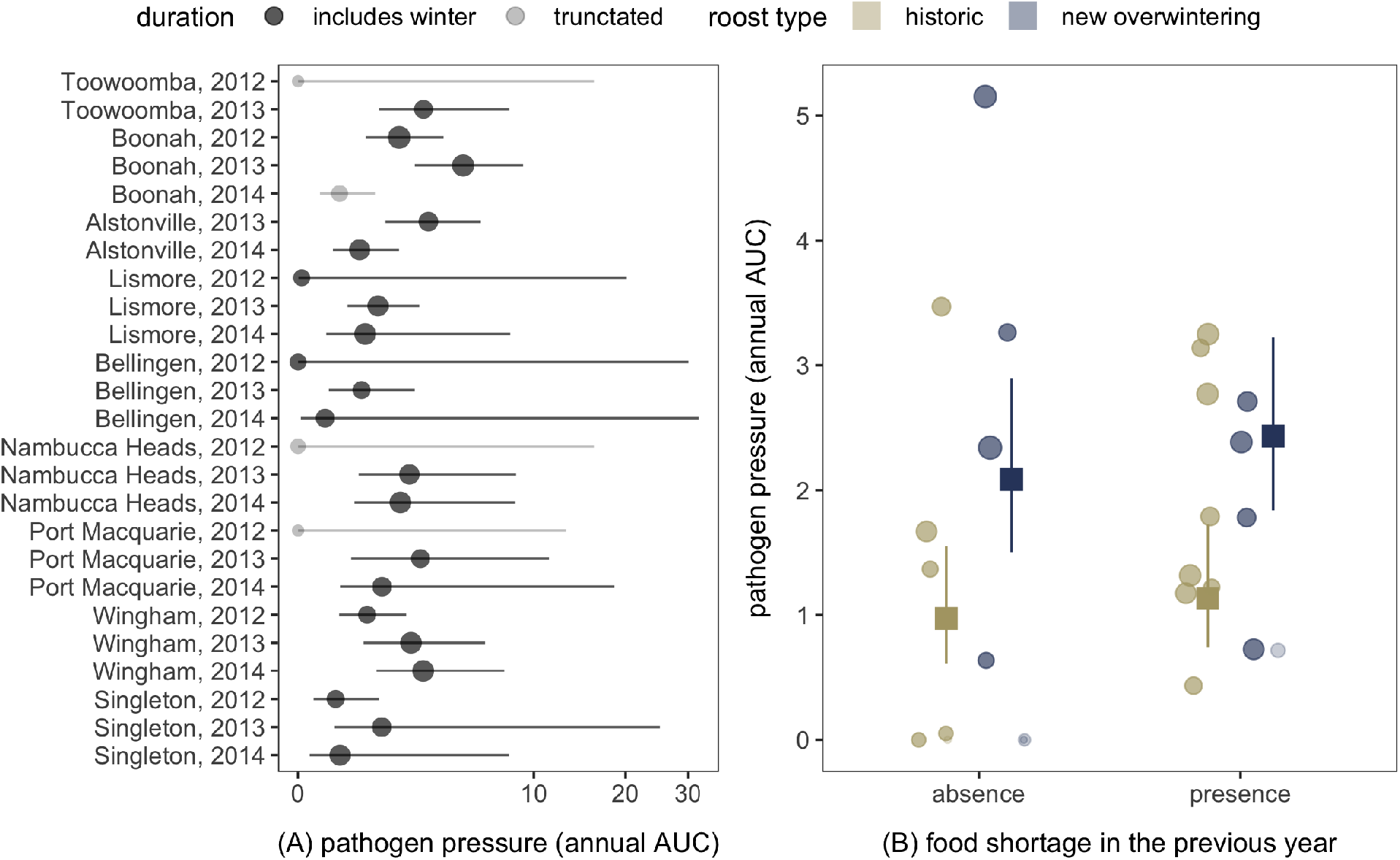
Variation in HeV pathogen pressure from flying foxes. The forest plot displays annual AUC estimates and 95% confidence intervals ordered by latitude and year; points are scaled by the number of sampling events per annual time series as weights. The horizontal axis uses a modulus transformation to accommodate wide upper bounds of some confidence intervals. Right displays fitted values and 95% confidence intervals for the GAM, with raw data (scaled by the number of sampling events per each time series) and modeled means colored by roost type. Transparency denotes AUC estimates derived from truncated annual time series (≤ 20 weeks).

### Pathogen pressure and HeV spillover

Within the period of flying fox sampling (2012–2014), 12 spillover events occurred within 500 km of our nine study roosts (Fig. 4A, Table S7). When aggregating spillovers within 50 km of each roost, our GAM explained 53% of the deviance and was primarily driven by spatial patterns, as AUC had a weak, nonlinear relationship with spillover (*χ^2^*=4.74, *p*=0.11). However, when expanding the buffers between 100 and 500 km from each roost, our GAMs explained 29– 45% of the deviance and predicted stronger associations between AUC and spillover (Table S8– S13). Generally, GAMs suggested a generally positive relationship between pathogen pressure and spillovers that was either maximized or saturated at intermediate-to-high AUC (Fig. 4B).

**Figure 4.**
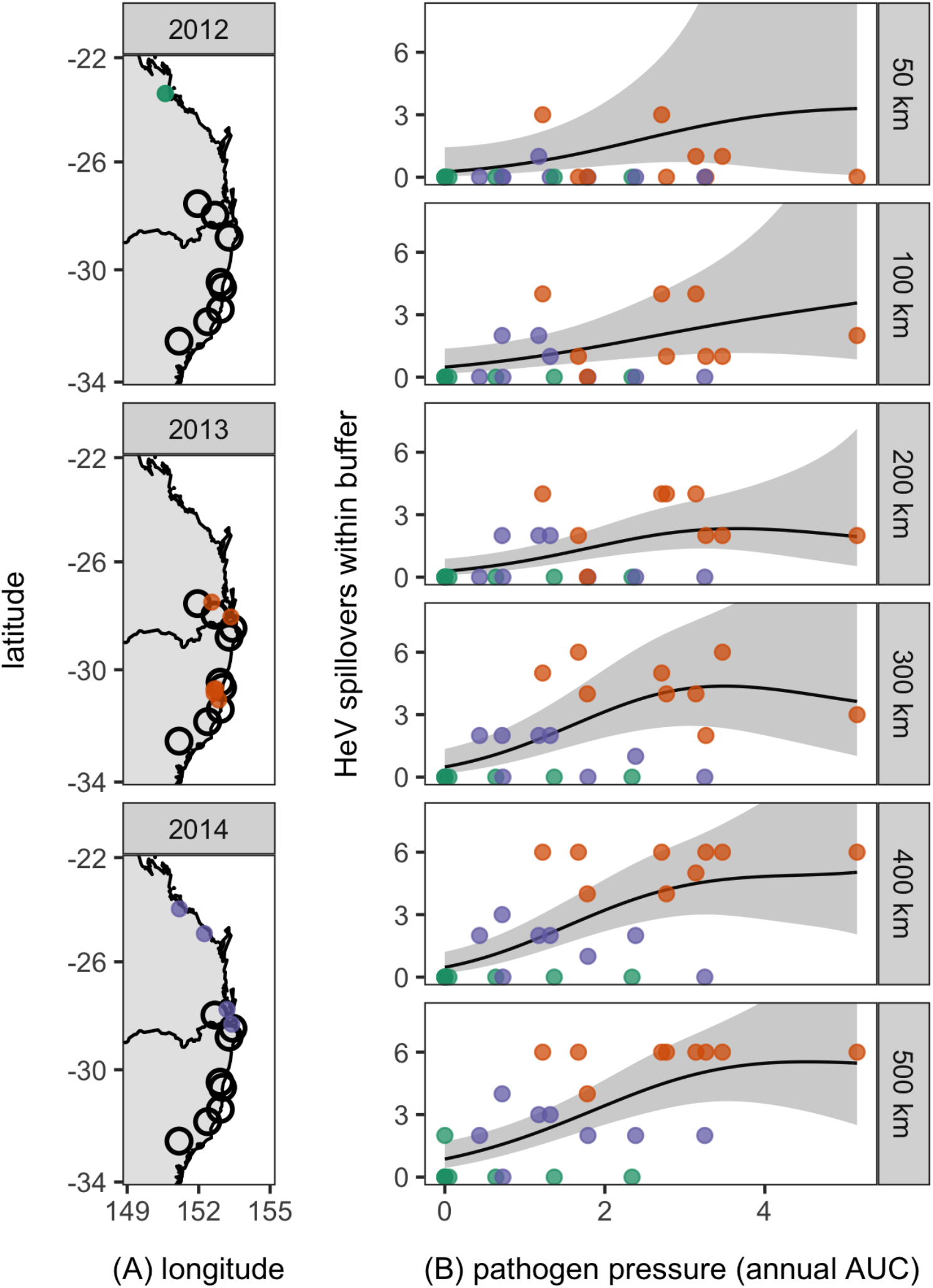
Spatiotemporal variation in regional HeV spillovers during the flying fox surveillance period (2012–2014) and its relationship with pathogen pressure (AUC). (A) Maps display the annual distributions of spillovers (colored by year) in relation to the nine analyzed roosts. Modeled relationships between AUC and spillover counts are shown with fitted values and 95% confidence intervals from GAMs for 50, 100, 200, 300, 400, and 500 km buffers of each roost.

## Discussion

Mechanistic understanding of cross-species transmission risk remains hindered by limited information on the relationships between reservoir host ecology, pathogen pressure, and observed spillover events [1,6,7,15]. By combining data on long-term ecology with the spatial and temporal distribution of infection, we here demonstrate that the ecological conditions experienced by reservoir hosts shape pathogen shedding. Prior work has shown that pulses of HeV excretion from flying foxes often occur in the Austral winter and that drier conditions in prior seasons are associated with peak annual shedding [20,21]. However, contextualizing these seasonal and climatic patterns with bat ecology here reveals that these periods of intensive virus shedding predominantly occur after food shortages and in new overwintering roosts in urban and agricultural habitats. By quantifying an aggregate metric of pathogen pressure, we also show that the accumulated intensity of HeV shedding provides distinct insights into these ecological drivers, as pathogen pressure is governed more by whether the roost is located in regions with native winter resources than by periods of recent food scarcity. Lastly, pathogen pressure was positively associated with cases of HeV in horses at a regional scale, which provides support for the spatiotemporal intensity of virus shedding as a mechanistic determinant of spillover risk.

Foremost, our findings emphasize that the ecological conditions experienced by reservoir hosts predict pronounced pulses of virus shedding and overall pathogen pressure. Across eastern Australia, habitats for the winter-flowering plants that drive flying fox nomadism have been reduced through land clearance, in turn driving nutritionally stressed bats into novel urban and agricultural environments outside their typical overwintering range (Eby et al., in review) [27,32,36,37]. Prior work has proposed, but not tested, that HeV shedding from flying foxes is driven by this process [23–25,63]. Acute food shortages could produce cumulative effects with other energetic and seasonal stressors (e.g., pregnancy, thermoregulation), as well as reliance on poor-quality, non-native food in agricultural and urban habitats, to alter within-host dynamics of viruses in bats [22,23]. Prior work has suggested that the physiological demands of winter thermoregulation in parts of the *P. alecto* range could drive observed negative correlations between minimum temperature and urinary cortisol excretion [40]. Our results support the idea of cumulative stressors, as HeV shedding was greatest not only in winter but also in roosts that were established in novel overwintering habitats and after a food shortage in the prior spring.

Increased viral shedding associated with suboptimal habitats or food shortages could be driven by immunosuppression, such that cumulative stressors may impair HeV tolerance and facilitate replication or allow latent infections to reactivate [25,64]. Recrudescent infection of henipaviruses in bats has been proposed to explain shedding pulses that coincide with stressors [24,25] and for seroconversions in captivity [65]. The mechanism driving the time lag between acute food shortages, which typically occur in spring, and increased HeV shedding in the following winter (i.e., in the subsequent 6–9 months; Fig. 1) remain unknown; however, the cumulative impact of multiple stressors (e.g., long-term nutritional stress, cold temperature, pregnancy) offers a potential explanation. Future studies, including immune assessments of wild bats and factorial experiments, will be critical to mechanistically understand the causal effect of spring food shortages, bat residency outside their winter range, dietary reliance on non-native plants, and other energetic stressors (e.g., pregnancy) on subsequent winter pulses of HeV shedding. Such work would also provide foundational insights into how bats control zoonotic viruses more broadly, given likely effects of stressors on immunity and tolerance [66,67].

Pathogen pressure has been suggested to be a critical determinant of spillover risk in recipient hosts [1,6], but quantifying the spatial and temporal distribution of shedding in ways that facilitate downstream analyses has remained challenging. Here, we propose the area under annual shedding curves (i.e., AUC) as a meaningful metric to summarize pathogen output into the environment. AUC has previously been used to summarize the cumulative pathogen output from within-host dynamics (e.g., *Mycoplasma agassizii* intensity across individual tortoises [68]) and number of infected individuals (e.g., fungal infections across *Daphnia* populations [69,70]), which capture host infectiousness and local epidemic size, respectively, rather than the explicit infectious dose available to recipient hosts. For the latter, epidemic size has in turn been linked to local ecological conditions, such as lake temperature and host community composition [69,70]; however, such metrics have not been applied to replicate reservoir host populations to identify predictors of pathogen pressure. In one case study of a zoonotic pathogen reservoir, data on avian influenza virus prevalence in ducks were used to derive AUC across three regions of North America, but inference on its underlying ecological drivers was limited by few replicates [53].

In our analyses, AUC not only displayed substantial heterogeneity, but also varied with roost ecology in ways that differed from seasonal GAMMs. Whereas the timing and amplitude of shedding was driven both by roost type and prior food shortages, AUC was exclusively driven by roost type. This contrast could relate to the tension between a short but intense shedding pulse and a smoldering shedding dynamic with lower intensity. Short but intense shedding pulses observed after food shortages could produce similar AUC to the lower amplitude shedding that occurred year-round in roosts without acute food scarcity. Such results emphasize how pathogen pressure can offer distinct insights into infection dynamics relative to shedding seasonality alone.

High pathogen pressure should theoretically have a pronounced effect on shaping the force of infection and the ultimate probability of spillover, given the hierarchical nature of cross-species transmission [1,6]. However, empirical support for pathogen pressure from reservoir hosts influencing disease cases in recipient hosts has been restricted by the rarity of spillover in many systems, such that evidence is mostly from temporal overlap between shedding events and outbreaks or broad-scale, regional analyses of reservoir infection and human disease data. For example, Marburg virus spillover to humans coincide with seasonal birth pulses of Egyptian fruit bats, when host shedding is most pronounced [71], and the density of bank voles seropositive for Puumala virus can also explain regional human incidence of nephropathia epidemica [72].

The relatively high frequency of HeV spillovers compared to less-frequent spillover in many other systems provides a tractable system for linking pathogen pressure and cross-species transmission [73]. Our analyses show that pathogen pressure (i.e., annual AUC) positively predicts observed spillovers within the broad, regional scales of flying fox movements (i.e., up to 500 km). As a caveat, no spillovers were recorded within typical nightly foraging ranges (i.e., 25 km) of the roosts monitored in our study region during the period of HeV surveillance [26,32]. Our findings therefore assume that the ecological conditions experienced by unsampled roosts in closer proximity to spillover events may be similar to those of our sampled roosts up to 500 km. As our patterns were consistent across a broad span of spatial scales relevant to flying fox mobility, our results suggest that the association between AUC and spillover are likely more affected by interannual variation in pathogen pressure, such as that driven by acute food shortages. Associations between pathogen pressure and spillovers were also mostly positive and monotonic but generally not exclusively linear. Importantly, our measure of pathogen pressure accumulates processes in the spillover hierarchy—including reservoir host distribution, reservoir host density, infection prevalence, and infection intensity—up to and including shedding into the environment [1]. Downstream factors such as HeV survival in the environment and recipient host distribution and susceptibility may moderate effects of high pathogen pressure on spillover risk [6,74]. Additionally, short but high amplitude pulses of virus shedding, such as those observed in the year following acute food shortages, may ultimately provide a more sufficient HeV dose to horses, thus resulting in roughly equivalent spillover risks from moderate and high AUC [1,75].

Because summarizing the spatiotemporal intensity of infection both provided distinct insights into ecological drivers of pathogen shedding and predicted spillover at broad regional scales, we suggest quantifying pathogen pressure would be particularly tractable and useful in systems where recipient host exposure occurs following pathogen release into the environment. Such systems include but are not limited to bat filoviruses and coronaviruses, avian influenza viruses, and helminths, protozoa, and fecal-oral bacteria of various wildlife [4,71,76,77]. Careful sampling designs for these environmentally shed pathogens, such as under-roost sampling methods for bats, could facilitate estimating pathogen pressure and, in turn, spillover risks [78].

In conclusion, we show that the ecological conditions experienced by bat reservoir hosts shape the timing, magnitude, and cumulative intensity of zoonotic virus shedding in ways that subsequently predict observed spillover events. Importantly, such inferences were only possible by integrating both spatiotemporal data on infection with long-term studies of host ecology and behavior, which demonstrates the importance of long-term, spatially replicated studies of reservoir hosts for linking environmental change and zoonotic pathogen dynamics. Despite a long-standing recognition of spatial and temporal scale in ecological research [79], replication across both axes remains challenging, particularly over relevant time intervals [80]. Spatiotemporal sampling is especially critical in the study of infectious disease [9,11], because pathogen shedding and transmission are both inherently spatial and temporal processes [63,81]. Connecting such data with changing ecology of wildlife further requires studies of abiotic and biotic correlates and host behavior and demography at similar or biologically meaningful spatial and temporal scales [7,14,82]. Here, the ecological conditions that predict HeV shedding from flying foxes were derived from behavioral data collected over approximately 25 years and from diverse sources [27,28,37] (Eby et al., in review). Although collecting this kind of ecological data will accordingly present logistical difficulties, the growth of national research networks, global community consortiums, and remote sensing, among other large-scale efforts, should facilitate similar approaches to link spatiotemporal data on both host ecology and infection [83]. In turn, careful attention to the ecology of reservoir hosts through long-term monitoring could facilitate improved early detection and preemptive management of zoonotic spillover events.

## Acknowledgements

Funding was provided by the DARPA PREEMPT program Cooperative Agreement D18AC00031, the U.S. National Science Foundation (DEB-1716698), and the USDA National Institute of Food and Agriculture (Hatch project 1015891). AJP was supported by an Australian Research Council DECRA fellowship (DE190100710). Data were collected through the National Hendra Virus Research Program and are provided by courtesy of the State of Queensland, Australia through the Department of Agriculture and Fisheries and the State of New South Wales through the Department of Primary Industries [20]. We acknowledge the Biripi, Gumbainggir, Widjabul Wia-bal, Wonnarua, and Yuggera Ugarapul people, who are the Traditional Custodians of the land upon which this work was conducted. We thank Hume Field, David Jordan, Peter Kirkland, and other co-contributors of the open access HeV dataset for data access and members of the Plowright lab for helpful feedback on previous versions of this manuscript.

## Author contributions

DJB, PE, AJP, and RKP designed the study; PE, AJP, WM, and RKP collected, collated, and managed data; DJB analyzed data; and DJB wrote the manuscript with feedback from coauthors.

## Data availability

All HeV shedding data and flying fox data used in this analysis are freely available from the Queensland government (CC BY 4.0; https://www.data.qld.gov.au/dataset/hev-infection-flying-foxes-eastern-australia) [20]. Ecological covariates of study roosts are provided in Table 1. Data on HeV spillovers within 500 km of our study roosts between 2012–2014 are provided to the nearest town in Table S4; finer-scale data are confidential as they identify individual properties.

## Code availability

Code for estimating AUC from each annual time series per roost will be provided in the Supplemental Material upon acceptance.

## References

1. Plowright RK, Parrish CR, McCallum H, Hudson PJ, Ko AI, Graham AL, Lloyd-Smith JO. 2017 Pathways to zoonotic spillover. Nat. Rev. Microbiol. 15, 502–510.

2. Becker DJ, Albery GF, Kessler MK, Lunn TJ, Falvo CA, Czirják GÁ, Martin LB, Plowright RK. 2020 Macroimmunology: The drivers and consequences of spatial patterns in wildlife immune defence. J. Anim. Ecol. 89, 972–995.

3. Plowright RK, Reaser JK, Locke H, Woodley SJ, Patz JA, Becker DJ, Oppler G, Hudson PJ, Tabor GM. 2021 Land use-induced spillover: a call to action to safeguard environmental, animal, and human health. Lancet Planet. Health 5, e237–e245. (doi:10.1016/S2542-5196(21)00031-0)

4. Ezenwa VO. 2004 Interactions among host diet, nutritional status and gastrointestinal parasite infection in wild bovids. Int. J. Parasitol. 34, 535–542.

5. Hernandez SM et al. 2016 Urbanized White Ibises (Eudocimus albus) as Carriers of Salmonella enterica of Significance to Public Health and Wildlife. PLOS ONE 11, e0164402. (doi:10.1371/journal.pone.0164402)

6. Washburne Alex D., Crowley Daniel E., Becker Daniel J., Manlove Kezia R., Childs Marissa L., Plowright Raina K. 2019 Percolation models of pathogen spillover. Philos. Trans. R. Soc. B Biol. Sci. 374, 20180331. (doi:10.1098/rstb.2018.0331)

7. Becker DJ, Washburne AD, Faust CL, Mordecai EA, Plowright RK. 2019 The problem of scale in the prediction and management of pathogen spillover. Philos. Trans. R. Soc. B Biol. Sci. 374, 20190224. (doi:10.1098/rstb.2019.0224)

8. Sokolow Susanne H. et al. 2019 Ecological interventions to prevent and manage zoonotic pathogen spillover. Philos. Trans. R. Soc. B Biol. Sci. 374, 20180342. (doi:10.1098/rstb.2018.0342)

9. Plowright RK, Becker DJ, McCallum H, Manlove KR. 2019 Sampling to elucidate the dynamics of infections in reservoir hosts. Philos. Trans. R. Soc. B (doi:10.1098/rstb.2018.0336)

10. Nusser SM, Clark WR, Otis DL, Huang L. 2008 Sampling considerations for disease surveillance in wildlife populations. J. Wildl. Manag. 72, 52–60.

11. Becker DJ, Crowley DE, Washburne AD, Plowright RK. 2019 Temporal and spatial limitations in global surveillance for bat filoviruses and henipaviruses. Biol. Lett. 15, 20190423. (doi:10.1098/rsbl.2019.0423)

12. Giron-Nava A et al. 2017 Quantitative argument for long-term ecological monitoring. Mar. Ecol. Prog. Ser. 572, 269–274.

13. Lindenmayer DB et al. 2012 Value of long-term ecological studies. Austral Ecol. 37, 745–757.

14. Ostfeld RS, Canham CD, Oggenfuss K, Winchcombe RJ, Keesing F. 2006 Climate, Deer, Rodents, and Acorns as Determinants of Variation in Lyme-Disease Risk. PLoS Biol. 4, e145. (doi:10.1371/journal.pbio.0040145)

15. Plowright RK et al. 2015 Ecological dynamics of emerging bat virus spillover. Proc R Soc B 282, 20142124. (doi:10.1098/rspb.2014.2124)

16. Murray K et al. 1995 A morbillivirus that caused fatal disease in horses and humans. Science 268, 94–97.

17. Eby P, Plowright R, McCallum H, Peel AJ. 2020 Conditions predict heightened Hendra virus spillover risk in horses this winter: actions now can change outcomes. Aust. Vet. J. 98, 270–271. (doi:https://doi.org/10.1111/avj.12964)

18. Halpin K et al. 2011 Pteropid bats are confirmed as the reservoir hosts of henipaviruses: a comprehensive experimental study of virus transmission. Am. J. Trop. Med. Hyg. 85, 946–951. (doi:10.4269/ajtmh.2011.10-0567)

19. Field H, Young P, Yob JM, Mills J, Hall L, Mackenzie J. 2001 The natural history of Hendra and Nipah viruses. Microbes Infect. 3, 307–314. (doi:10.1016/S1286-4579(01)01384-3)

20. Field H et al. 2015 Spatiotemporal Aspects of Hendra Virus Infection in Pteropid Bats (Flying-Foxes) in Eastern Australia. PLOS ONE 10, e0144055. (doi:10.1371/journal.pone.0144055)

21. Páez DJ, Giles J, McCallum H, Field H, Jordan D, Peel AJ, Plowright RK. 2017 Conditions affecting the timing and magnitude of Hendra virus shedding across pteropodid bat populations in Australia. Epidemiol. Infect., 1–11. (doi:10.1017/S0950268817002138)

22. Plowright RK, Foley P, Field HE, Dobson AP, Foley JE, Eby P, Daszak P. 2011 Urban habituation, ecological connectivity and epidemic dampening: the emergence of Hendra virus from flying foxes (Pteropus spp.). Proc. R. Soc. B Biol. Sci. 278, 3703–3712. (doi:10.1098/rspb.2011.0522)

23. Kessler MK et al. 2018 Changing resource landscapes and spillover of henipaviruses. Ann. N. Y. Acad. Sci. 1429, 78–99. (doi:10.1111/nyas.13910)

24. Peel AJ et al. 2019 Synchronous shedding of multiple bat paramyxoviruses coincides with peak periods of Hendra virus spillover. Emerg. Microbes Infect. 8, 1314–1323. (doi:10.1080/22221751.2019.1661217)

25. Plowright RK, Peel AJ, Streicker DG, Gilbert AT, McCallum H, Wood J, Baker ML, Restif O. 2016 Transmission or Within-Host Dynamics Driving Pulses of Zoonotic Viruses in Reservoir–Host Populations. PLoS Negl. Trop. Dis. 10, e0004796. (doi:10.1371/journal.pntd.0004796)

26. Eby P. 1991 Seasonal movements of grey-headed flying-foxes, Pteropus poliocephalus (Chiroptera: Pteropodidae), from two maternity camps in northern New South Wales. Wildl. Res. 18, 547–559.

27. Eby P, Richards G, Collins L, Parry-Jones K. 1999 The distribution, abundance and vulnerability to population reduction of a nomadic nectarivore, the Grey-headed Flying-fox Pteropus poliocephalus in New South Wales, during a period of resource concentration. Aust. Zool. 31, 240–253. (doi:10.7882/AZ.1999.024)

28. Eby P, Law B. 2008 Ranking the feeding habitat of grey-headed flying foxes for conservation management. Canberra Dep. Environ. Herit. Water Arts

29. Wilson BA, Neldner VJ, Accad A. 2002 The extent and status of remnant vegetation in Queensland and its implications for statewide vegetation management and legislation. Rangel. J. 24, 6–35.

30. Giles JR, Plowright RK, Eby P, Peel AJ, McCallum H. 2016 Models of Eucalypt phenology predict bat population flux. Ecol. Evol. 6, 7230–7245.

31. Plowright RK, Field HE, Smith C, Divljan A, Palmer C, Tabor G, Daszak P, Foley JE. 2008 Reproduction and nutritional stress are risk factors for Hendra virus infection in little red flying foxes (Pteropus scapulatus). Proc. R. Soc. B 275, 861–869. (doi:10.1098/rspb.2007.1260)

32. Giles JR, Eby P, Parry H, Peel AJ, Plowright RK, Westcott DA, McCallum H. 2018 Environmental drivers of spatiotemporal foraging intensity in fruit bats and implications for Hendra virus ecology. Sci. Rep. 8, 9555. (doi:10.1038/s41598-018-27859-3)

33. Páez DJ, Restif O, Eby P, Plowright RK. 2018 Optimal foraging in seasonal environments: implications for residency of Australian flying foxes in food-subsidized urban landscapes. Phil Trans R Soc B 373, 20170097. (doi:10.1098/rstb.2017.0097)

34. Markus N, Hall L. 2004 Foraging behaviour of the black flying-fox (Pteropus alecto) in the urban landscape of Brisbane, Queensland. Wildl. Res. 31, 345–355.

35. Mcdonald-Madden E, Schreiber ESG, Forsyth DM, Choquenot D, Clancy TF. 2005 Factors affecting Grey-headed Flying-fox (Pteropus poliocephalus: Pteropodidae) foraging in the Melbourne metropolitan area, Australia. Austral Ecol. 30, 600–608. (doi:10.1111/j.1442-9993.2005.01492.x)

36. Tait J, Perotto-Baldivieso HL, McKeown A, Westcott DA. 2014 Are Flying-Foxes Coming to Town? Urbanisation of the Spectacled Flying-Fox (Pteropus conspicillatus) in Australia. PLOS ONE 9, e109810. (doi:10.1371/journal.pone.0109810)

37. Lunn T, Eby P, Brooks R, McCallum H, Plowright R, Kessler M, Peel A. 2021 Conventional wisdom on roosting behaviour of Australian flying foxes-a critical review, and evaluation using new data. Authorea Prepr.

38. Van Der Ree R, McDonnell MJ, Temby I, Nelson J, Whittingham E. 2006 The establishment and dynamics of a recently established urban camp of flying foxes (Pteropus poliocephalus) outside their geographic range. J. Zool. 268, 177–185.

39. Williams NSG, Mcdonnell MJ, Phelan GK, Keim LD, Van Der Ree R. 2006 Range expansion due to urbanization: Increased food resources attract Grey-headed Flying-foxes (Pteropus poliocephalus) to Melbourne. Austral Ecol. 31, 190–198. (doi:10.1111/j.1442-9993.2006.01590.x)

40. McMichael L, Edson D, Smith C, Mayer D, Smith I, Kopp S, Meers J, Field H. 2017 Physiological stress and Hendra virus in flying-foxes (Pteropus spp.), Australia. PLOS ONE 12, e0182171. (doi:10.1371/journal.pone.0182171)

41. Wang H-H, Kung NY, Grant WE, Scanlan JC, Field HE. 2013 Recrudescent infection supports hendra virus persistence in Australian flying-fox populations. PloS One 8, e80430.

42. Edson D et al. 2019 Time of year, age class and body condition predict Hendra virus infection in Australian black flying foxes (Pteropus alecto). Epidemiol. Infect. 147.

43. Boardman WSJ, Baker ML, Boyd V, Crameri G, Peck GR, Reardon T, Smith IG, Caraguel CGB, Prowse TAA. 2020 Seroprevalence of three paramyxoviruses; Hendra virus, Tioman virus, Cedar virus and a rhabdovirus, Australian bat lyssavirus, in a range expanding fruit bat, the Grey-headed flying fox (Pteropus poliocephalus). PLOS ONE 15, e0232339. (doi:10.1371/journal.pone.0232339)

44. Peel A, Eby P, Kessler M, Lunn T, Breed A, Plowright R. 2017 Hendra virus spillover risk in horses: heightened vigilance and precautions being urged this winter. Aust. Vet. J. 95, N20–N21.

45. Smith C, Skelly C, Kung N, Roberts B, Field H. 2014 Flying-fox species density-a spatial risk factor for Hendra virus infection in horses in Eastern Australia. PLoS One 9, e99965.

46. Goldspink LK, Edson DW, Vidgen ME, Bingham J, Field HE, Smith CS. 2015 Natural Hendra virus infection in flying-foxes-tissue tropism and risk factors. PLoS One 10, e0128835.

47. Edson D et al. 2015 Routes of Hendra Virus Excretion in Naturally-Infected Flying-Foxes: Implications for Viral Transmission and Spillover Risk. PLOS ONE 10, e0140670. (doi:10.1371/journal.pone.0140670)

48. McKee CD, Islam A, Luby SP, Salje H, Hudson PJ, Plowright RK, Gurley ES. 2021 The Ecology of Nipah Virus in Bangladesh: A Nexus of Land-Use Change and Opportunistic Feeding Behavior in Bats. Viruses 13, 169.

49. Edson D, Field H, McMichael L, Jordan D, Kung N, Mayer D, Smith C. 2015 Flying-Fox Roost Disturbance and Hendra Virus Spillover Risk. PLOS ONE 10, e0125881. (doi:10.1371/journal.pone.0125881)

50. Smith IL, Halpin K, Warrilow D, Smith GA. 2001 Development of a fluorogenic RT-PCR assay (TaqMan) for the detection of Hendra virus. J. Virol. Methods 98, 33–40.

51. Wood S. 2006 Generalized additive models: an introduction with R. CRC press.

52. R Core Team. 2013 R: A language and environment for statistical computing. R Foundation for Statistical Computing. Vienna, Austria. See http://www.R-project.org.

53. Lisovski S, Hoye BJ, Klaassen M. 2017 Geographic variation in seasonality and its influence on the dynamics of an infectious disease. Oikos 126, 931–936. (doi:10.1111/oik.03796)

54. Cosgrove CL, Wood MJ, Day KP, Sheldon BC. 2008 Seasonal variation in Plasmodium prevalence in a population of blue tits Cyanistes caeruleus. J. Anim. Ecol. 77, 540–548.

55. Wood SN, Augustin NH. 2002 GAMs with integrated model selection using penalized regression splines and applications to environmental modelling. Ecol. Model. 157, 157–177. (doi:10.1016/S0304-3800(02)00193-X)

56. Pedersen EJ, Miller DL, Simpson GL, Ross N. 2019 Hierarchical generalized additive models in ecology: an introduction with mgcv. PeerJ 7, e6876.

57. Burnham KP, Anderson DR. 2002 Model selection and multimodel inference: a practical information-theoretic approach. Springer Science & Business Media.

58. Bolstad WM. 2010 Understanding computational Bayesian statistics. John Wiley & Sons.

59. Viechtbauer W. 2010 Conducting meta-analyses in R with the metafor package. J. Stat. Softw. 36, 1–48.

60. Senior AM, Grueber CE, Kamiya T, Lagisz M, O’Dwyer K, Santos ESA, Nakagawa S. 2016 Heterogeneity in ecological and evolutionary meta-analyses: its magnitude and implications. Ecology 97, 3293–3299. (doi:10.1002/ecy.1591)

61. Roberts BJ, Catterall CP, Eby P, Kanowski J. 2012 Long-distance and frequent movements of the flying-fox Pteropus poliocephalus: implications for management. PLoS One 7, e42532.

62. Kilinc BK, Asfha HD. 2019 Penalized Splines Fitting for a Poisson Response Including Outliers. Pak. J. Stat. Oper. Res., 979–988.

63. Lunn TJ, Peel AJ, McCallum H, Eby P, Kessler MK, Plowright RK, Restif O. 2021 Spatial dynamics of pathogen transmission in communally roosting species: impacts of changing habitats on bat-virus dynamics. J. Anim. Ecol. n/a. (doi:10.1111/1365-2656.13566)

64. Virgin HW, Wherry EJ, Ahmed R. 2009 Redefining chronic viral infection. Cell 138, 30–50. (doi:10.1016/j.cell.2009.06.036)

65. Glennon EE et al. 2019 What is stirring in the reservoir? Modelling mechanisms of henipavirus circulation in fruit bat hosts. Philos. Trans. R. Soc. B Biol. Sci. 374, 20190021. (doi:10.1098/rstb.2019.0021)

66. Subudhi S, Rapin N, Misra V. 2019 Immune System Modulation and Viral Persistence in Bats: Understanding Viral Spillover. Viruses 11, 192. (doi:10.3390/v11020192)

67. Banerjee A, Baker ML, Kulcsar K, Misra V, Plowright R, Mossman K. 2020 Novel insights into immune systems of bats. Front. Immunol. 11, 26.

68. Aiello CM, Esque TC, Nussear KE, Emblidge PG, Hudson PJ. 2019 The slow dynamics of mycoplasma infections in a tortoise host reveal heterogeneity pertinent to pathogen transmission and monitoring. Epidemiol. Infect. 147.

69. Penczykowski RM, Hall SR, Civitello DJ, Duffy MA. 2014 Habitat structure and ecological drivers of disease. Limnol. Oceanogr. 59, 340–348.

70. Shocket MS, Strauss AT, Hite JL, Šljivar M, Civitello DJ, Duffy MA, Cáceres CE, Hall SR. 2018 Temperature drives epidemics in a zooplankton-fungus disease system: a trait-driven approach points to transmission via host foraging. Am. Nat. 191, 435–451.

71. Amman BR et al. 2012 Seasonal Pulses of Marburg Virus Circulation in Juvenile Rousettus aegyptiacus Bats Coincide with Periods of Increased Risk of Human Infection. Plos Pathog. 8. (doi:10.1371/journal.ppat.1002877)

72. Tersago K, Verhagen R, Vapalahti O, Heyman P, Ducoffre G, Leirs H. 2011 Hantavirus outbreak in Western Europe: reservoir host infection dynamics related to human disease patterns. Epidemiol. Infect. 139, 381–390. (doi:10.1017/S0950268810000956)

73. Plowright RK, Hudson PJ. 2021 From Protein to Pandemic: The Transdisciplinary Approach Needed to Prevent Spillover and the Next Pandemic. Viruses 13, 1298. (doi:10.3390/v13071298)

74. Childs ML, Nova N, Colvin J, Mordecai EA. 2019 Mosquito and primate ecology predict human risk of yellow fever virus spillover in Brazil. Philos. Trans. R. Soc. B 374, 20180335.

75. Lunn TJ, Restif O, Peel AJ, Munster VJ, De Wit E, Sokolow S, Van Doremalen N, Hudson P, McCallum H. 2019 Dose–response and transmission: the nexus between reservoir hosts, environment and recipient hosts. Philos. Trans. R. Soc. B 374, 20190016.

76. Becker Daniel J. et al. 2018 Assessing the contributions of intraspecific and environmental sources of infection in urban wildlife: Salmonella enterica and white ibis as a case study. J. R. Soc. Interface 15, 20180654. (doi:10.1098/rsif.2018.0654)

77. Breban R, Drake JM, Stallknecht DE, Rohani P. 2009 The Role of Environmental Transmission in Recurrent Avian Influenza Epidemics. PLOS Comput. Biol. 5, e1000346. (doi:10.1371/journal.pcbi.1000346)

78. Giles JR, Peel AJ, Wells K, Plowright RK, McCallum H, Restif O. 2018 Optimizing non-invasive sampling of an infectious bat virus. bioRxiv, 401968. (doi:10.1101/401968)

79. Levin SA. 1992 The problem of pattern and scale in ecology: the Robert H. MacArthur award lecture. Ecology 73, 1943–1967.

80. Estes L, Elsen PR, Treuer T, Ahmed L, Caylor K, Chang J, Choi JJ, Ellis EC. 2018 The spatial and temporal domains of modern ecology. Nat. Ecol. Evol. 2, 819–826. (doi:10.1038/s41559-018-0524-4)

81. Grenfell BT, Bjørnstad ON, Kappey J. 2001 Travelling waves and spatial hierarchies in measles epidemics. Nature 414, 716–723. (doi:10.1038/414716a)

82. Lunn T, Peel AJ, Eby P, Brooks R, Plowright RK, Kessler MK, Mccallum H. In press. Counterintuitive scaling between population size and density: implications for modelling transmission of infectious diseases in bat populations.

83. Dietze MC et al. 2018 Iterative near-term ecological forecasting: Needs, opportunities, and challenges. Proc. Natl. Acad. Sci. 115, 1424–1432.

