## Supplemental Material for "Ecological conditions experienced by bat reservoir hosts predict the intensity of Hendra virus excretion over space and time"

Figure S1. Relative abundance of *Pteropus alecto* from weekly total flying fox counts across each of the nine time series. Red lines indicate the maximum fraction of *P. alecto* per roost. We used the 25% maximum occupancy from Singleton as our cutoff for this bat species being common or rare in GAMMs, as this was the lowest roost-level maximum fraction of this species.

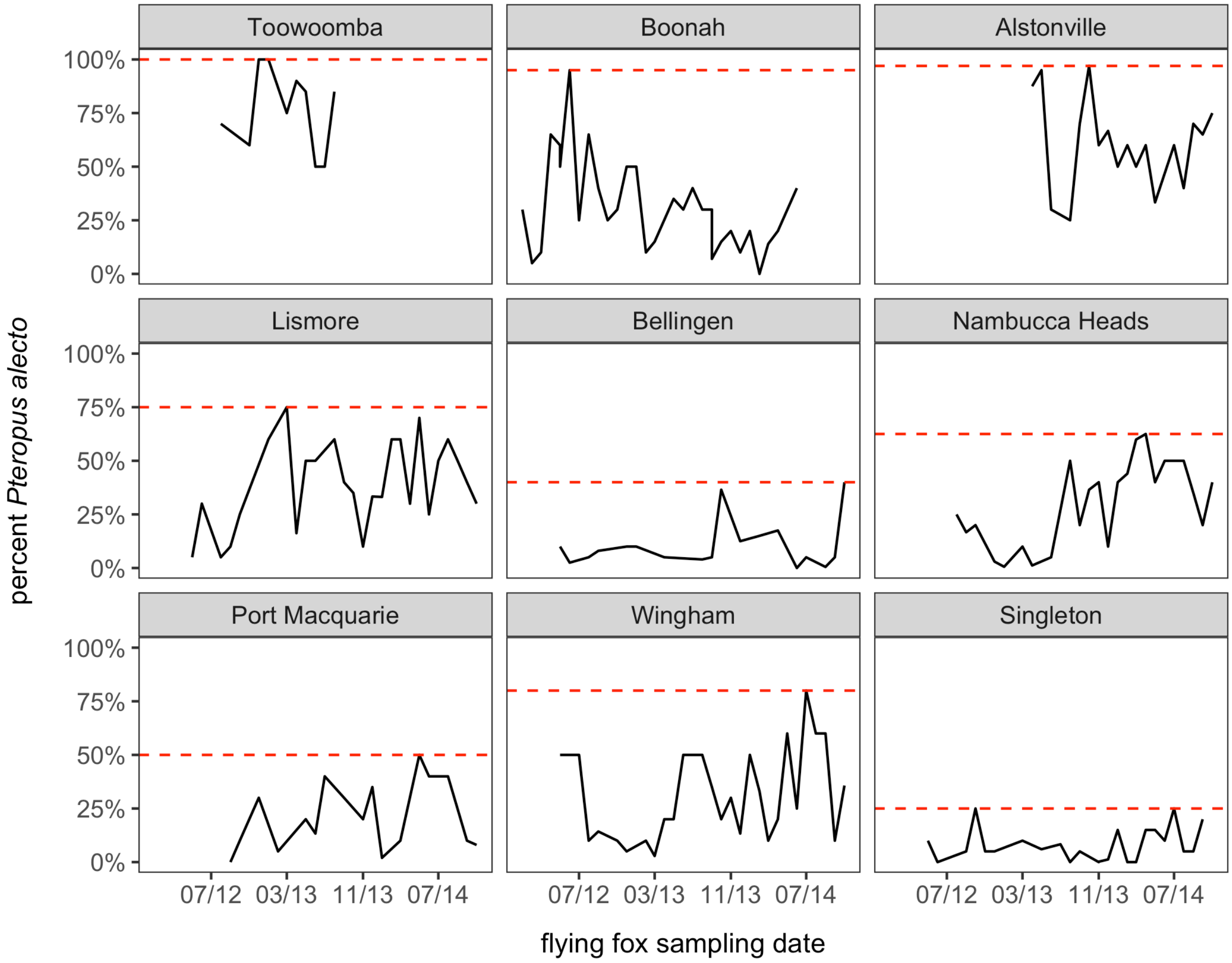

Figure S2. Fitted values from GAMs of each roost per year. GAMs included a cyclic cubic spline for week except for two roost–years with uniformly zero prevalence and sparse time series (i.e., Toowoomba and Port Macquarie 2012), which used thin plate splines for convergence. Models were fit with REML, and we integrated fitted values to derive AUC. Raw data are overlaid.

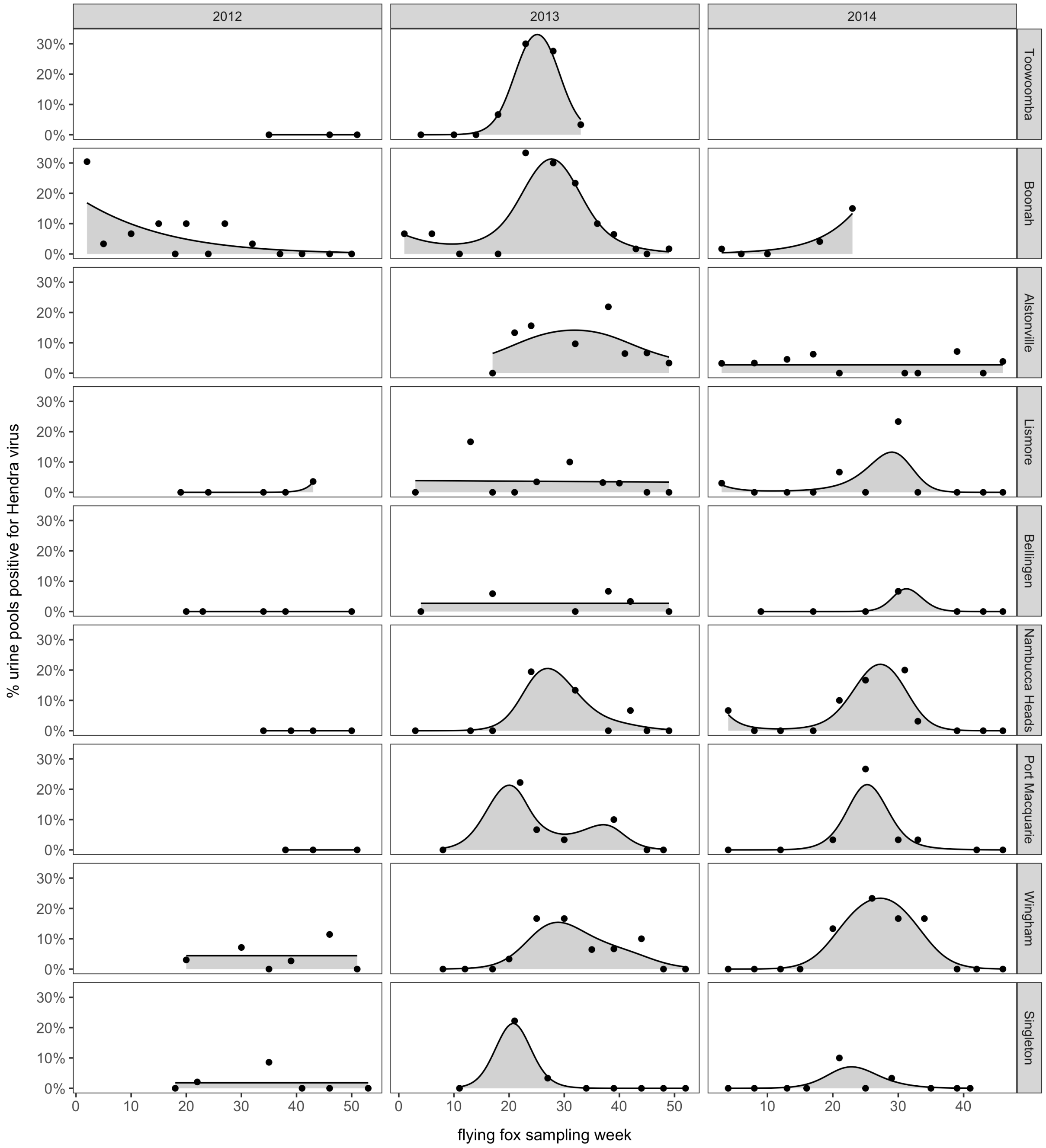

Figure S3. Modeled relationship between the mean annual (*n*=25), median annual (*n*=25), and maximum winter (*n*=24) proportion of *Pteropus alecto* per roost and HeV pathogen pressure (annual AUC). Lines and bands display fitted values and 95% confidence intervals for each GAM, with raw data overlaid  and scaled by the number of sampling events per each time series.

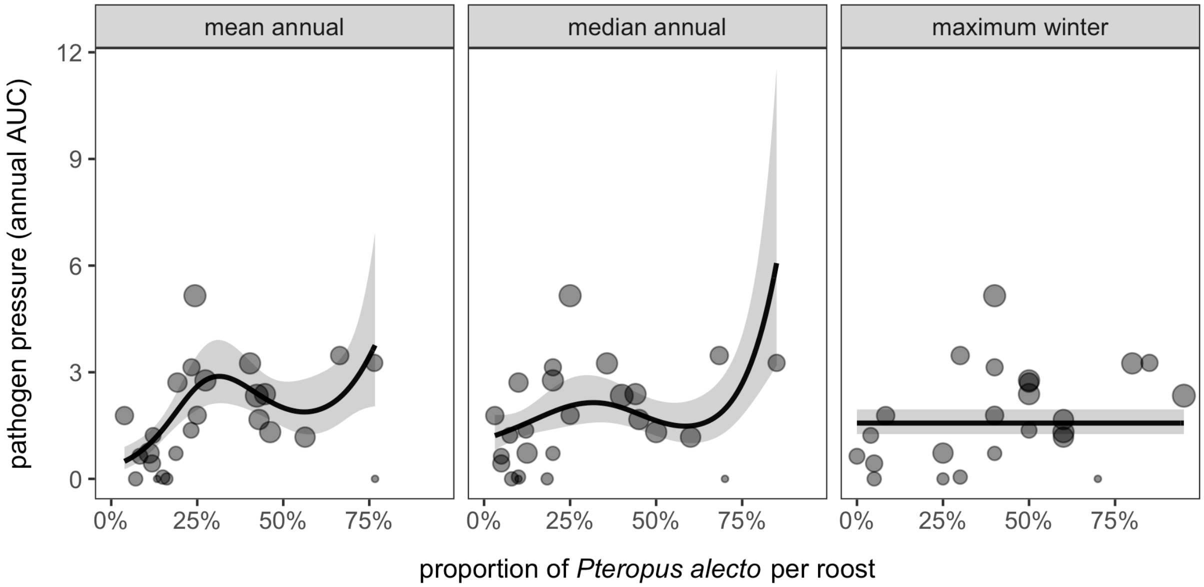

Figure S4. Modeled relationship between ecological predictors of annual AUC after excluding four estimates from truncated annual time series (i.e., 20 or fewer sampled weeks). Shown are the fitted values and 95% confidence intervals for each GAM overlaid with raw data (scaled by the number of sampling events per each time series) and modeled means colored by roost type.

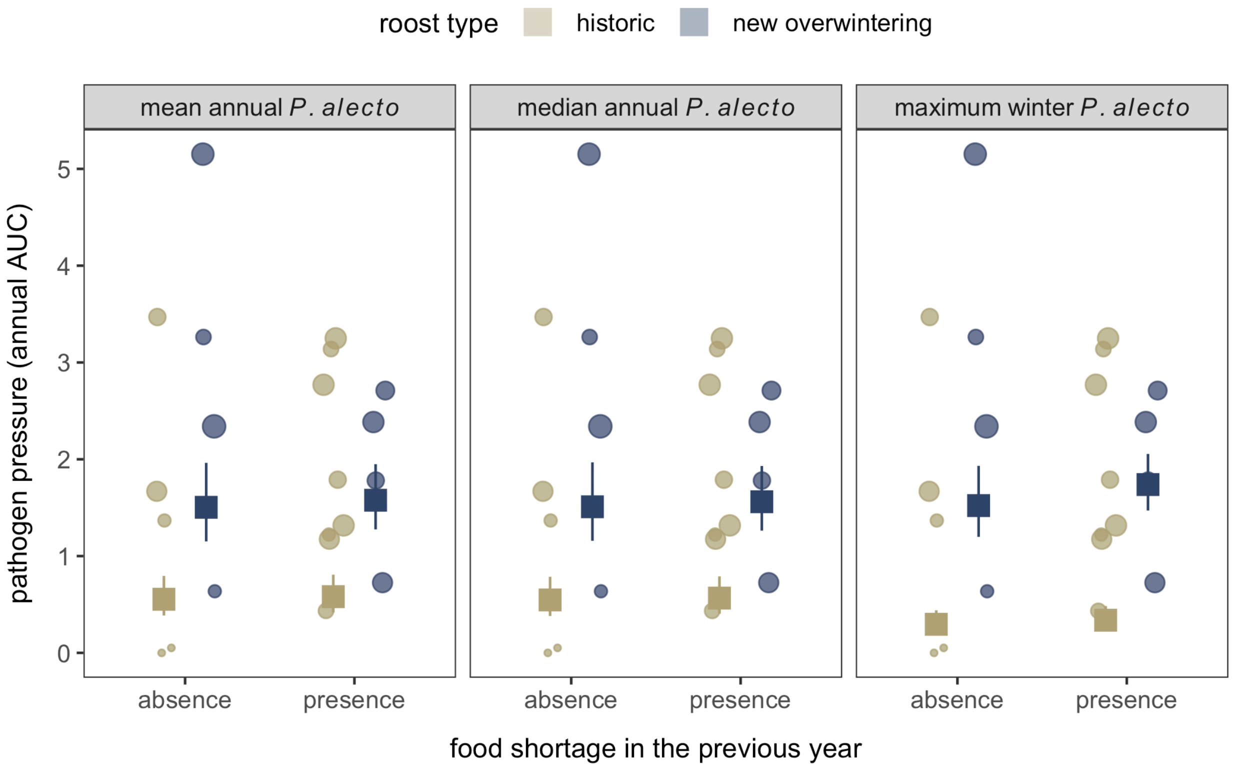

Table S1. Nested GAMMs predicting seasonal HeV shedding from Australian flying foxes. Competing models are ranked by ΔAICc with the Akaike weights ($w_i$), adjusted $R^2$, and the proportion of deviance explained. All GAMMs include a bivariate smooth of longitude and latitude to control for spatial dependence and random factor smooth of week per roost per year.

| Fixed effects | ΔAICc | $w_i$ | $R^2$ | % deviance |
|---|---|---|---|---|
| ~ shortage + type + s(week) + s(week, by=shortage) + s(week, by=type) | 0.00 | 0.59 | 0.66 | 0.71 |
| ~ shortage * type + s(week) + s(week, by=shortage) + s(week, by=type) | 0.85 | 0.39 | 0.65 | 0.71 |
| ~ shortage * type + s(week) + s(week, by=shortage) + s(week, by=type) + s(week, by=(shortage, type)) | 7.31 | 0.02 | 0.65 | 0.71 |

Table S2. Results from the GAM of AUC estimates from roosts using median annual proportions of *Pteropus alecto* (*n*=25), fit using REML. Fixed effects are presented as ordered factors with model coefficients (categorical) or the estimated degrees of freedom (EDF) and test statistics.

| Term | *β* | *t* | EDF | *F* | *p* |
|---|---|---|---|---|---|
| Intercept | -0.11 | -0.80 | | | 0.43 |
| New overwintering | 1.11 | 5.64 | | | <0.001 |
| Prior food shortage | 0.16 | 1.50 | | | 0.16 |
| s(median percent *Pteropus alecto*) | | | 3.44 | 5.56 | <0.001 |
| s(longitude, latitude) | | | 3.83 | 7.67 | <0.001 |

Table S3. Results from the GAM of AUC estimates from roosts using maximum winter proportions of *P. alecto* (*n*=24; one roost–year had no winter sampling), fit using REML. Fixed effects are presented as ordered factors with model coefficients or the EDF and test statistics.

| Term | *β* | *t* | EDF | *F* | *p* |
|---|---|---|---|---|---|
| Intercept | 0.09 | 0.85 | | | 0.41 |
| New overwintering | 0.89 | 5.99 | | | <0.001 |
| Prior food shortage | 0.07 | 0.65 | | | 0.52 |
| s(maximum winter percent *Pteropus alecto*) | | | 0.00 | 0.00 | 0.77 |
| s(longitude, latitude) | | | 3.92 | 24.82 | <0.001 |

Table S4. Results from the GAM of AUC excluding four sparse annual time series (≤20 weeks) using mean annual proportions of *Pteropus alecto* (*n*=21), fit using REML. Fixed effects are presented as ordered factors with model coefficients (categorical) or EDF and test statistics.

| Term | *β* | *t* | EDF | *F* | *p* |
|---|---|---|---|---|---|
| Intercept | -0.16 | -1.55 | | | 0.15 |
| New overwintering | 1.34 | 8.56 | | | <0.001 |
| Prior food shortage | 0.25 | 2.62 | | | 0.02 |
| s(median percent *Pteropus alecto*) | | | 3.56 | 7. 65 | <0.001 |
| s(longitude, latitude) | | | 3.89 | 13.90 | <0.001 |

Table S5. Results from the GAM of AUC excluding four sparse annual time series (≤20 weeks) using median annual proportions of *Pteropus alecto* (*n*=21), fit using REML. Fixed effects are presented as ordered factors with model coefficients (categorical) or EDF and test statistics.

| Term | *β* | *t* | EDF | *F* | *p* |
|---|---|---|---|---|---|
| Intercept | 0.03 | 0.38 | | | 0.71 |
| New overwintering | 1.07 | 8.99 | | | <0.001 |
| Prior food shortage | 0.13 | 1.29 | | | 0.22 |
| s(maximum winter percent *Pteropus alecto*) | | | 0.88 | 1.31 | 0.03 |
| s(longitude, latitude) | | | 3.95 | 43.43 | <0.001 |

Table S6. Results from the GAM of AUC excluding four sparse annual time series (≤20 weeks) using maximum winter proportions of *Pteropus alecto* (*n*=21), fit using REML. Fixed effects are presented as ordered factors with model coefficients (categorical) or EDF and test statistics.

| Term | *β* | *t* | EDF | *F* | *p* |
|---|---|---|---|---|---|
| Intercept | -0.18 | -1.99 | | | 0.07 |
| New overwintering | 1.64 | 10.79 | | | <0.001 |
| Prior food shortage | 0.13 | 1.46 | | | 0.17 |
| s(maximum winter percent *Pteropus alecto*) | | | 3.88 | 16.10 | 0.77 |
| s(longitude, latitude) | | | 3.96 | 43.78 | <0.001 |

Table S7. Year and location of the 12 HeV spillovers within 500 km of our nine study roosts between 2012–2014. Finer-scale data are confidential as they identify individual properties.

| Nearest regional center | state | year |
|---|---|---|
| Rockhampton | Queensland | 2012 |
| Rockhampton | Queensland | 2012 |
| Macksville | New South Wales | 2013 |
| Brisbane Valley | Queensland | 2013 |
| Macksville | New South Wales | 2013 |
| Gold Coast | Queensland | 2013 |
| Kempsey | New South Wales | 2013 |
| Kempsey | New South Wales | 2013 |
| Bundaberg | Queensland | 2014 |
| Beenleigh | Queensland | 2014 |
| Murwillumbah | New South Wales | 2014 |
| Calliope | Queensland | 2014 |

Table S8. GAM of HeV spillovers within 50 km of each roost per year, fit using REML. Fixed effects are presented with their EDF and test statistics. The GAM explained 53% of the deviance.

| Term | EDF | $\chi^2$ | *p* |
|---|---|---|---|
| s(pathogen pressure) | 1.51 | 4.73 | 0.11 |
| s(longitude, latitude) | 2.62 | 7.55 | 0.03 |

Table S9. GAM of HeV spillovers within 100 km of each roost per year, fit using REML. Fixed effects are presented with their EDF and test statistics. The GAM explained 33% of the deviance.

| Term | EDF | $\chi^2$ | *p* |
|---|---|---|---|
| s(pathogen pressure) | 1.55 | 7.87 | 0.03 |
| s(longitude, latitude) | 2.16 | 4.67 | 0.08 |

Table S10. GAM of HeV spillovers within 200 km of each roost per year, fit using REML. Fixed effects are presented with their EDF and test statistics. The GAM explained 29% of the deviance.

| Term | EDF | $\chi^2$ | *p* |
|---|---|---|---|
| s(pathogen pressure) | 1.96 | 9.68 | 0.01 |
| s(longitude, latitude) | 0.61 | 0.84 | 0.23 |

Table S11. GAM of HeV spillovers within 300 km of each roost per year, fit using REML. Fixed effects are presented with their EDF and test statistics. The GAM explained 42% of the deviance.

| Term | EDF | $\chi^2$ | *p* |
|---|---|---|---|
| s(pathogen pressure) | 2.17 | 14.63 | <0.01 |
| s(longitude, latitude) | 2.37 | 6.00 | 0.05 |

Table S12. GAM of HeV spillovers within 400 km of each roost per year, fit using REML. Fixed effects are presented with their EDF and test statistics. The GAM explained 45% of the deviance.

| Term | EDF | $\chi^2$ | *p* |
|---|---|---|---|
| s(pathogen pressure) | 2.33 | 19.98 | <0.01 |
| s(longitude, latitude) | 1.20 | 2.86 | 0.08 |

Table S13. GAM of HeV spillovers within 500 km of each roost per year, fit using REML. Fixed effects are presented with their EDF and test statistics. The GAM explained 41% of the deviance.

| Term | EDF | $\chi^2$ | *p* |
|---|---|---|---|
| s(pathogen pressure) | 2.00 | 19.83 | <0.01 |
| s(longitude, latitude) | 0.71 | 1.36 | 0.14 |
